# Sequence and ionic requirements of pUG fold quadruplexes

**DOI:** 10.1101/2025.10.28.685102

**Authors:** Saeed Roschdi, Takuma Kume, Riley J. Petersen, Abby McCann, Cristian A. Escobar, Anika Richard, Samuel E. Butcher

**Affiliations:** Department of Biochemistry, University of Wisconsin-Madison, Madison, WI, USA

## Abstract

Poly(UG) or “pUG” repeat RNA can fold into a left-handed parallel quadruplex, the pUG fold. The pUG fold directs the amplification of RNAi in *C. elegans*, and pUG sequences are abundant in eukaryotic transcriptomes. Here, we report the sequence and ionic requirements for pUG folding. The pUG fold requires 12 guanosines but has an otherwise flexible sequence requirement. The uridines can be substituted with other nucleotides, with some sequence variants folding better than pUG RNA. The GA repeat sequence (GA)_12_ also forms a pUG-like fold, albeit with lower thermodynamic stability than (GU)_12_. The pUG fold can tolerate multiple deoxyribose substitutions but does not fold when the backbone is entirely deoxyribose. It has a high affinity and specificity for potassium ions (K_1/2_ = 6 mM) and does not fold in sodium or ammonium ions. Addition of 2 mM Mg^2+^ does not further stabilize the pUG fold, and the polyamines spermine and spermidine slightly decrease its stability. Finally, the pUG fold is sensitive to surrounding sequence context and complementary flanking sequences can stabilize pUG folds, while other sequences can interfere with folding. These data expand our understanding of RNA folding and provide a basis for understanding and predicting pUG folds.

## Introduction

Dinucleotide repeats are common elements in eukaryotic transcriptomes, with poly(UG) or “pUG” repeats being among the most abundant (1). We previously described the pUG fold, a unique left-handed parallel intramolecular quadruplex (G4) that induces gene silencing by directing RNAi amplification in *C. elegans* (1,2). The shortest sequence that can form a pUG fold is the 23-nucleotide (nt) sequence (GU)_11.5_ (1). This sequence is extraordinarily abundant in human RNAs, occurring over 20,000 times. The polymorphic expansion of genomic GT repeats encoding pUG sequences has been associated with diseases including cancer and cystic fibrosis (1). We determined the crystal and solution structures of the 24-nt RNA (GU)_12_, which forms a pUG fold (1,3)(Figure 1). The pUG fold has three G quartets, one U quartet and 3 centrally coordinated potassium ions (Figure 1). Additionally, the pUG fold has 4 bulged uridines, 3 single uridine propeller loops and, in the case of (GU)_12_, a flexible 3’ terminal uridine that is not required for the fold (1)(Figure 1). The pUG fold is distinct from other known quadruplexes as it contains no consecutive guanosines in its sequence and is the only known example of an RNA G4 with an overall left-handed topology. The left-handed topology is due to Z-form *syn*-*anti* backbone inversions within the center of the fold (1,3,4). The 24-nt pUG fold structure has pseudo-4-fold symmetry, with 4 repeating units of a reverse S-shaped hexamer GUGUGU (Figure 1). Each nucleotide of the hexamer is in a different conformation with respect to glycosidic torsion angle, sugar pucker, and backbone orientation (Figure 1).

**Figure 1.**
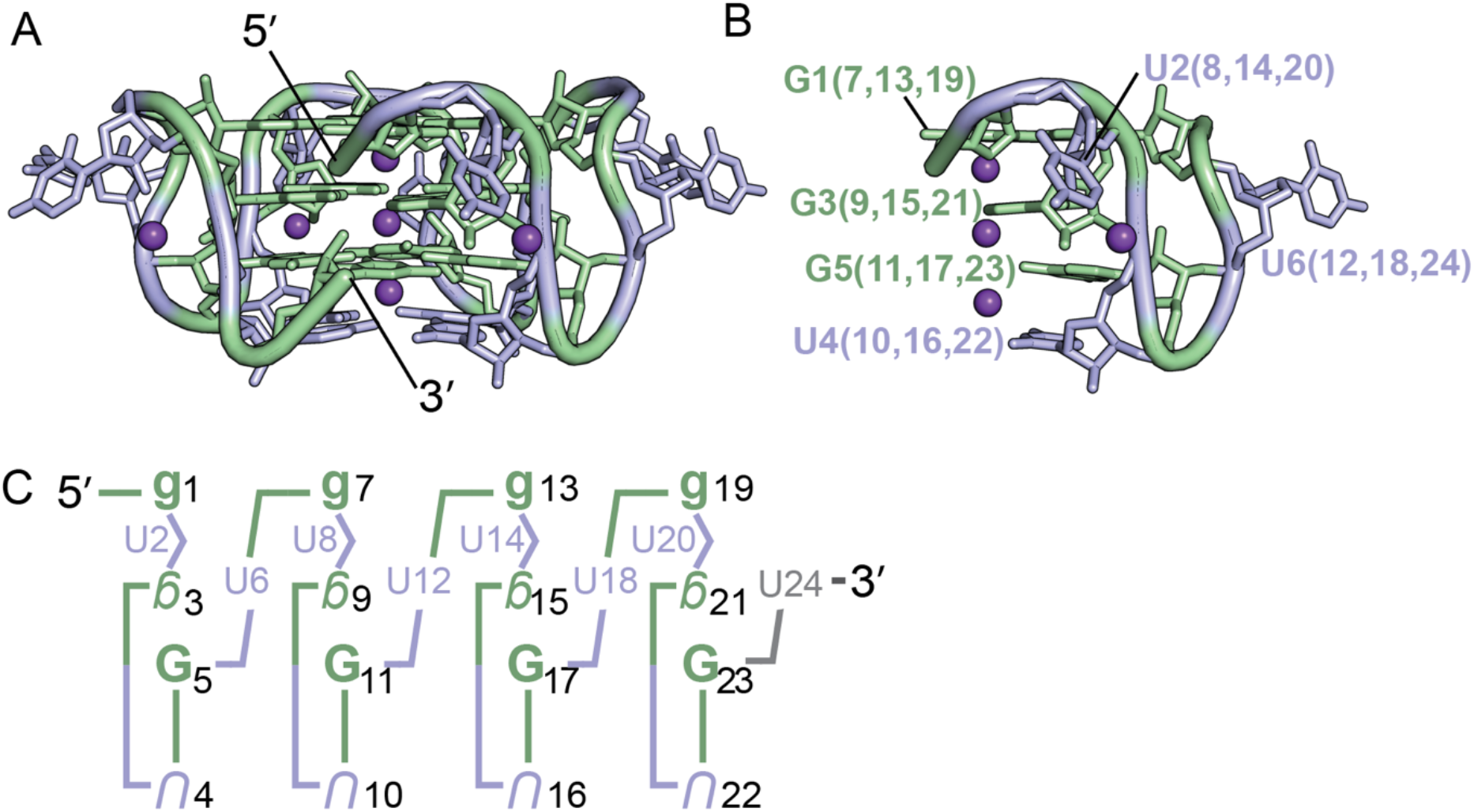
(A) Structure of the (GU)_12_ pUG fold (PDB: 7MKT). Guanosines are shown in green, uridines in blue, and potassium ions in purple (shown as one-fourth reduced size for clarity) (B) Close-up view of one-fourth of the molecule, showing the reverse S-shaped backbone of the GUGUGU hexamer, a conformation that is repeated four times. (C) Schematic of the pUG fold. Upper case is *anti*, lower case is *syn*. Backbone inversions are indicated by upside down letters. C3’ endo is italicized, C2’ endo is bold. The last 3’ uridine (grey) is disordered.

RNA structures can be challenging to predict, especially for non-A-form tertiary structures (5). For example, the pUG fold is a stable structure that forms both *in vitro* and *in vivo* (1) but is not predicted by AlphaFold 3 (6). RNA structure prediction is complicated by the fact that even a perfectly matched sequence-structure relationship does not necessarily provide evidence for a fold. This is particularly true of flexible RNAs that are best described as ensembles of structures, the distribution of which can be influenced by the process of transcription, surrounding RNA sequence, ions, pH, and the availability of small molecule ligands and protein partners (7-9). Regarding pUG RNA, an interesting example is found in fish, which have thousands of perfect matches to the pUG fold sequence within introns. However, these pUGs are also accompanied by complementary CA repeats, which form long-range CA/UG base-pairing interactions that bring together the 5’ and 3’ ends of introns and replace the need for the splicing factor U2AF2 (10). On the other hand, human RNAs contain thousands of pUG repeat sequences in introns which are not accompanied by complementary CA repeats regions (1). Humans have at least one protein that can specifically interact with pUG folds (11).

The sequence requirements and cation specificity of the pUG fold have not been described. We have previously shown that the pUG fold has some flexibility as it can tolerate AA insertions (1) and single deoxynucleotide substitutions (3). G4 structures typically coordinate potassium ions, but some can also fold by coordinating sodium or ammonium ions (12-14). The polyamines spermine and spermidine are also present at micromolar concentrations in human cells and can either stabilize or destabilize G4s (15-18). It has been proposed that some RNA G4 structures may sense and regulate polyamine levels in cells (19).

Here we report the sequence and ionic requirements for pUG fold formation. We show that 12 guanosines are strictly required for the fold, but the uridines can be individually substituted by any nucleotide. Multiple U to A substitutions are allowed, and even the sequence (GA)_12_ forms a pUG-like fold. The pUG fold requires low millimolar concentrations of potassium and cannot fold in buffers containing sodium or ammonium ions. The presence of Mg^2+^ does not stabilize the fold, and high micromolar concentrations of polyamines decrease its stability. The pUG fold can tolerate multiple deoxyribose substitutions but cannot be entirely deoxyribose. Finally, we show that surrounding sequences can either stabilize or interfere with pUG folding.

## Results

### Single nucleotide variants of the pUG fold

To investigate the sequence requirements of the pUG fold, we created all possible single nucleotide variations within one of the reverse S-shaped hexameric repeats of the pUG fold (Figure 1B) and assessed folding via circular dichroism (CD) spectroscopy (Figure 2A and B). CD reports on the different base stacking geometries within the fold. The negative 244 and positive 260 nm peaks are typical of right-handed *anti-anti* stacking in parallel quadruplexes, which is observed at (U4-G5). The positive 280 nm peak corresponds to the unusual *syn-syn* stacking observed at G1-G3, and the negative 304 nm peak corresponds to the left-handed (Z-form) *syn-anti* stacking of the G3 and G5 quartets (20). We targeted the second hexameric repeat of the pUG fold for mutagenesis, which allowed all of the RNA variants to be efficiently transcribed *in vitro* by T7 RNA polymerase. Analysis of CD spectra for the 19 different RNAs in 150 mM KCl buffer showed that none of the G substitutions are properly folded (Figure 2A). In contrast to the G variants, all U variants are well-folded and in fact, some fold slightly better than the reference sequence (GU)_12_ (Figure 2B). The intensity of the well-resolved negative CD peak at 304 nm provides an accurate and quantitative measure of the fraction of pUG folds in RNA, as it reports on the unique, left-handed conformation in the center of the fold (1,20). The fraction folded and standard deviation were determined from the CD measurements of 2 independently prepared samples (biological replicates). The most well-folded RNA is the sequence variant U12A (Figure 2B). In the pUG fold, U12 is unpaired and forms a single nucleotide propeller loop that spans four quartets (Figure 1). Another well-folded sequence variant is U8G, which occurs at the bulged-out conformation within the structure (Figure 2B). Finally, U10 variants are folded (Figure 2B) and this is surprising, as this position corresponds to the U quartet. These data suggest that the U quartet can be disrupted or is flexible enough to incorporate other nucleotides. Mixed A-U-A-U quartets have been previously observed (21). The average fraction folded for all RNAs, as determined by CD, is listed in Table 1.

**Table 1.**

| Sequence | Average Fraction Folded | Standard Deviation |
| --- | --- | --- |
| (GU)12 | 0.75 | 0.14 |
| G7A | 0.03 | 0.03 |
| G7C | 0.05 | 0.01 |
| G7U | 0.08 | 0.02 |
| U8A | 0.75 | 0.02 |
| U8C | 0.76 | 0.12 |
| U8G | 0.89 | 0.11 |
| G9A | 0.00 | 0.03 |
| G9C | 0.04 | 0.03 |
| G9U | 0.04 | 0.03 |
| U10A | 0.95 | 0.09 |
| U10C | 0.75 | 0.03 |
| U10G | 0.73 | 0.24 |
| G11A | 0.02 | 0.04 |
| G11C | 0.11 | 0.03 |
| G11U | 0.05 | 0.02 |
| U12A | 1.00 | 0.00 |
| U12C | 0.81 | 0.05 |
| U12G | 0.61 | 0.02 |
| U2A | 0.86 | 0.03 |
| U4A | 0.90 | 0.10 |
| U6A | 0.93 | 0.09 |
| U2A, U8A, U14A, U20A | 0.85 | 0.05 |
| U4A, U10A, U16A, U22A | 0.75 | 0.01 |
| U6A, U12A, U18A, U24A | 0.78 | 0.11 |
| (GA)12 | 0.67 | 0.01 |
| Random tails | 0.06 | 0.01 |
| AU bps | 0.59 | 0.04 |
| Mixed bps | 0.59 | 0.01 |
| Mango bps | 0.50 | 0.03 |
| d(GT)12 | 0.00 | 0.01 |
| d(GU)12 | 0.00 | 0.01 |
| dT2, dT8, dT16, dT20 | 0.80 | 0.01 |
| dG3, dG9, dG15, dG21 | 0.60 | 0.01 |
| d(T2,G3)x4 | 0.64 | 0.02 |
| d(G1, T2, G3)x4 | 0.50 | 0.01 |
| d(G1, T2, G5)x4 | 0.01 | 0.04 |
| d(G1, T2, T6)x4 | 0.47 | 0.02 |
| d(G1, G5, T6)x4 | 0.52 | 0.01 |
| d(T2, G5, T6)x4 | 0.00 | 0.04 |
| d(T4, G5, T6)x4 | 0.43 | 0.03 |
| d(G1, T2, G5, T6)x4 | 0.00 | 0.01 |
| d(G1, U2, G5, U6)x4 | 0.31 | 0.03 |
| d(G1, T4, G5, T6)x4 | 0.12 | 0.01 |

**Figure 2.**
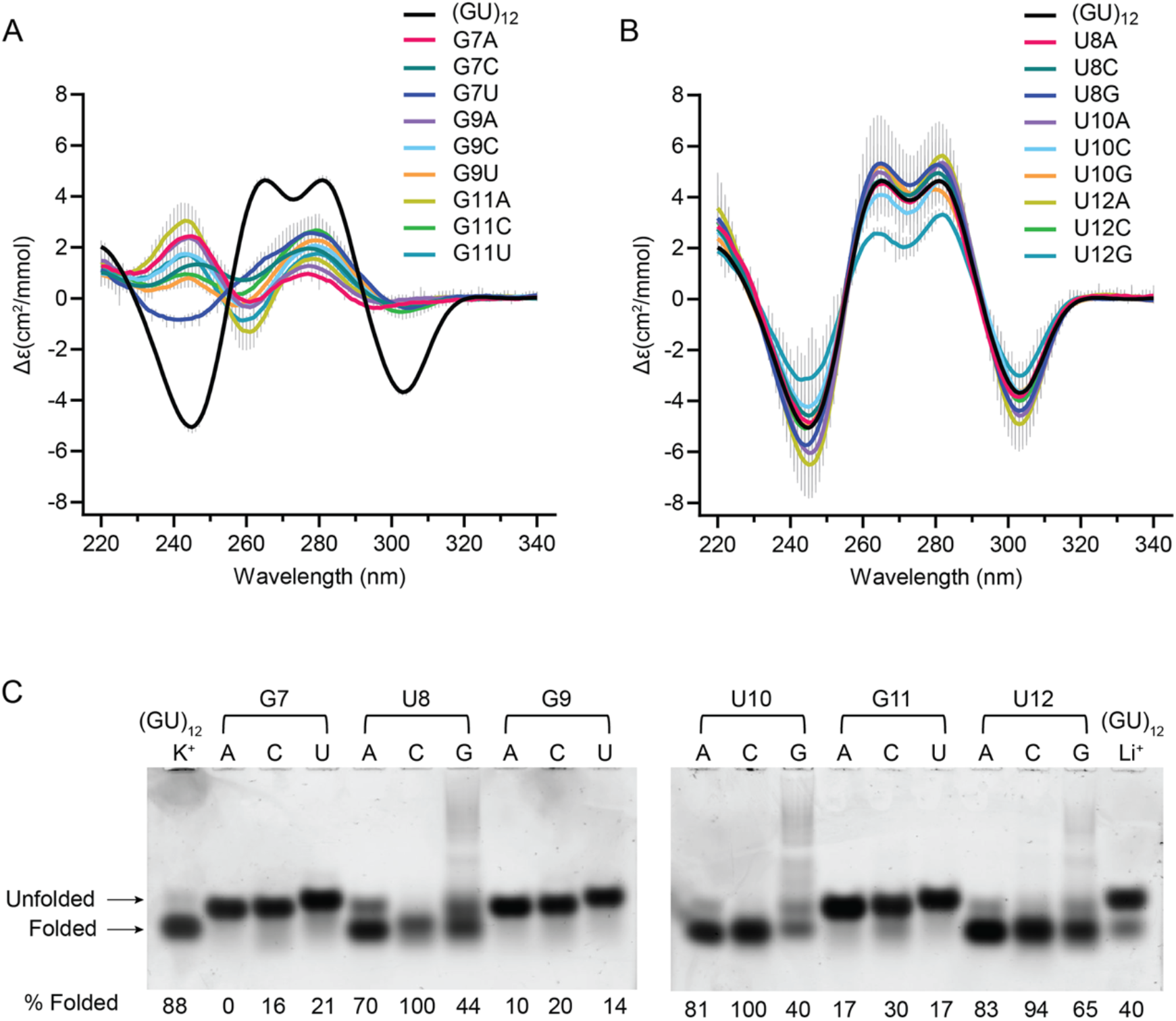
Circular dichroism spectra of single nucleotide variants that are (A) unfolded (B) folded. Error bars from duplicate measurements of biological replicates are shown with vertical gray lines. (C) Native gel electrophoresis of all single nucleotide variants for positions 7-12. First and last lanes contain (GU)_12_ in K^+^ and Li^+^, respectively. Apparent percent folded is listed at the bottom of the gel.

We further analyzed the sequence variants by native gel electrophoresis, in which 5 mM KCl was included in the gel and running buffer to maintain folding. In this experiment, the (GU)_12_ control RNA has a fraction folded of 0.88, which agrees with the CD data (0.75 + 0.14)(Figure 2 and Table 1) and prior NMR-based measurements (20). An unfolded (GU)_12_ control RNA in 150 mM LiCl partially re-folds during electrophoresis in the potassium gel and provides markers for the electrophoretic mobilities of the folded and unfolded forms of the RNA (Figure 2C, last lane). The EMSA data confirm that the G variants are unfolded, and the U variants are folded (Figure 2C). An exception for the G variants is G11C, which appears to retain a small fraction of folded molecules. This agrees with the CD data showing a small negative peak for G11C corresponding to a folded fraction of 0.11 (Figure 2A and Table 1). These data suggest that a small fraction of molecules can incorporate a cytidine within the central G5-11-17-23 quartet of the molecule. Mixed G-C-G-C quartets have been previously observed in G4 structures (22). The EMSA assay shows the U to A or C variants are well-folded, while the U to G variants are partially folded (Figure 2C), and the CD measurements show that U12G is the least well folded of the U variants (Figure 2B). The U to G variants display smearing in the gel, which is not observed for the other variants (Figure 2C). As these U to G variants create GGG stretches, we hypothesize the smearing is due to heterogenous intermolecular G4 interactions.

Since the 4-fold symmetry of the pUG fold is only broken by the 5’ and 3’ ends, it seems likely that sequence variations at symmetry-related positions should have similar effects on folding. To check this, we analyzed U to A variants within the first hexameric repeat (U2A, U4A and U6A)(Figure 3) and compared these variants to the symmetry-related sequence variants in the second hexameric repeat (U8A, U10A and U12A)(Figure 2). Indeed, the resulting data are similar and within the standard deviation of the measurements for all but U2A and U8A. The U2A, U4A and U6A have folded fractions of 0.86, 0.90 and 0.93, respectively, while the symmetry-related variants U8A, U10A and U12A have folded fractions of 0.75, 0.95 and 1.0, respectively (Table 1). We further analyzed the thermodynamic stabilities of the U to A variants (U2A, U4A, and U6A)(Figure 3). Temperature melting data were fit to the modified Gibbs-Helmholtz equation to determine the T_m_ (23). The reference sequence (GU)_12_ has a T_m_ = 52 °C (1). The U2A and U6A variants have slightly decreased thermodynamic stabilities, with T_m_ = 49 °C (Figure 3 B,D). In contrast a large decrease in stability is observed for the U-quartet variant U4A, with T_m_ = 44 °C (Figure 3C). Therefore, although substitutions in the U quartet are tolerated, the U quartet adds stability to the overall fold.

**Figure 3.**
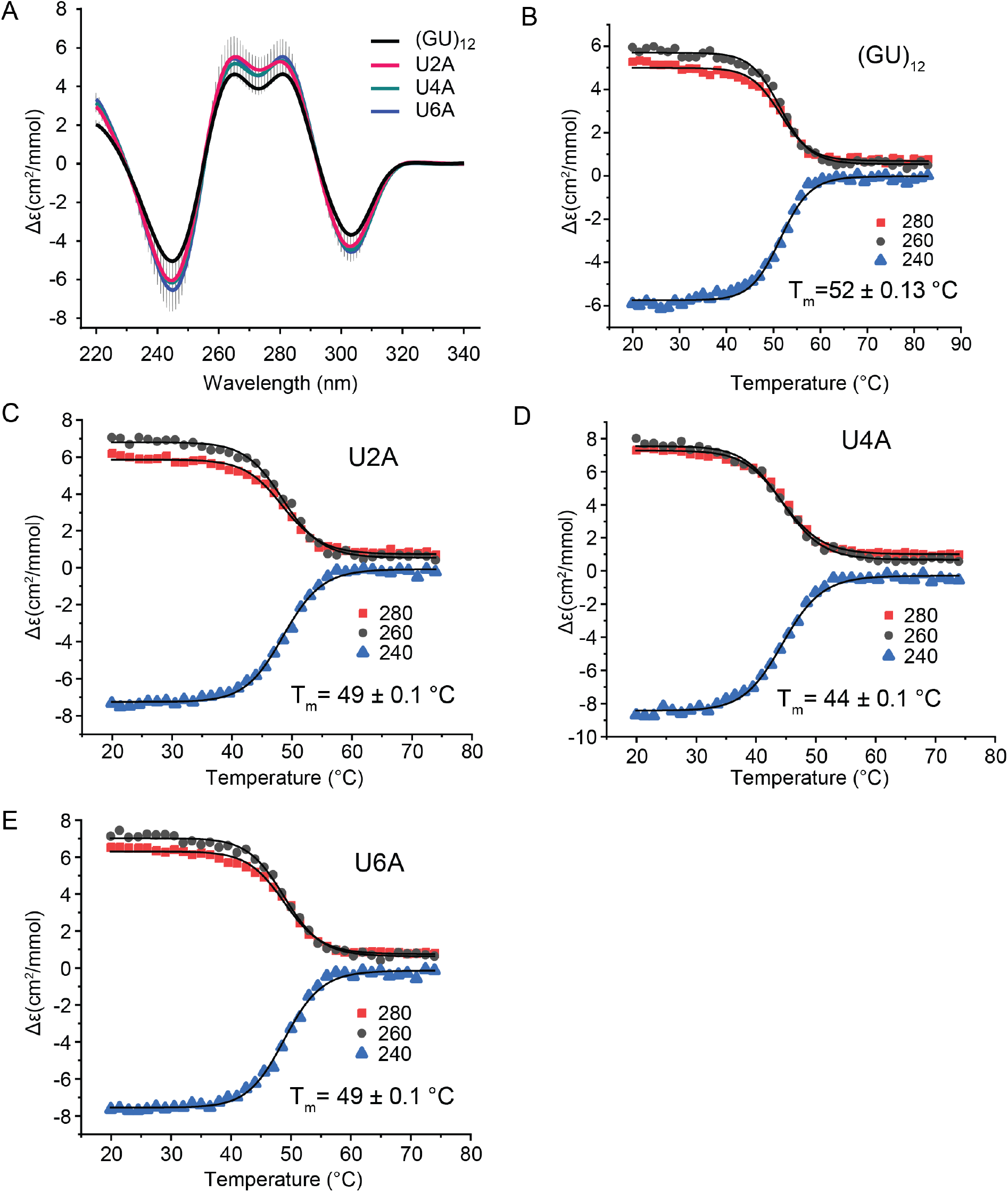
(A) Comparison of CD spectra of (GU)_12_ and U2A, U4A and U6A sequence variants. CD monitored thermal denaturation of (B) (GU)_12_, (C) U2A variant, (D) U4A variant, and (E) U6A variant measured at three wavelengths. All data fitted to modified Gibbs-Helmholtz equation to determine T_m_.

### Extensive sequence variants of the pUG fold

To further investigate the sequence flexibility of the pUG fold, we measured the folding of sequence variants containing multiple U to A substitutions (Figure 4A). These variants included quadruple U to A substitutions, and the sequence (GA)_12_. All RNAs were folded and display the negative 304 nm peak characteristic of the Z-form *syn-anti* stacking of the central G3 and G5 quartets (Figure 4A). The quadruple bulged nucleotide variant (U2A, U8A, U14A, U20A) has a much smaller 280 nm peak but is otherwise folded (Figure 4A). The 280 nm peak corresponds to the *syn-syn* stacking conformation of the G1-7-13-19 and G3-9-15-21 quartets, and the (U2A, U8A, U14A, U20A) bulge variants occur between these quartets. Therefore, adenosines at this bulge position may alter the stacking geometry of the G1 and G3 quartets while leaving the rest of the fold intact. The quadruple U to A variant at the U quartet (U4A, U10A, U16A, U22A) is folded and displays all 4 peaks associated with the pUG fold (Figure 4A). These data further demonstrate that the U quartet is not necessary for pUG folding. This is likely because the U4 quartet is located at a solvent exposed edge of the structure and only forms 4 hydrogen bonds, vs. the G quartets which have 8 hydrogen bonds (1,3). The quadruple U to A propeller loop variant (U6A, U12A, U18A, U24A) is also well-folded (Figure 4A and Table 1). These data demonstrate that the pUG fold can tolerate multiple U to A substitutions, at any position in the fold. We therefore asked if all uridines can be replaced by adenosine and tested the folding of (GA)_12_. Similar to the bulged variant (U2A, U8A, U14A, U20A), the (GA)_12_ variant displays a pUG-like fold that is missing the 280 nm peak but is otherwise folded (Figure 4A and Table 1).

**Figure 4.**
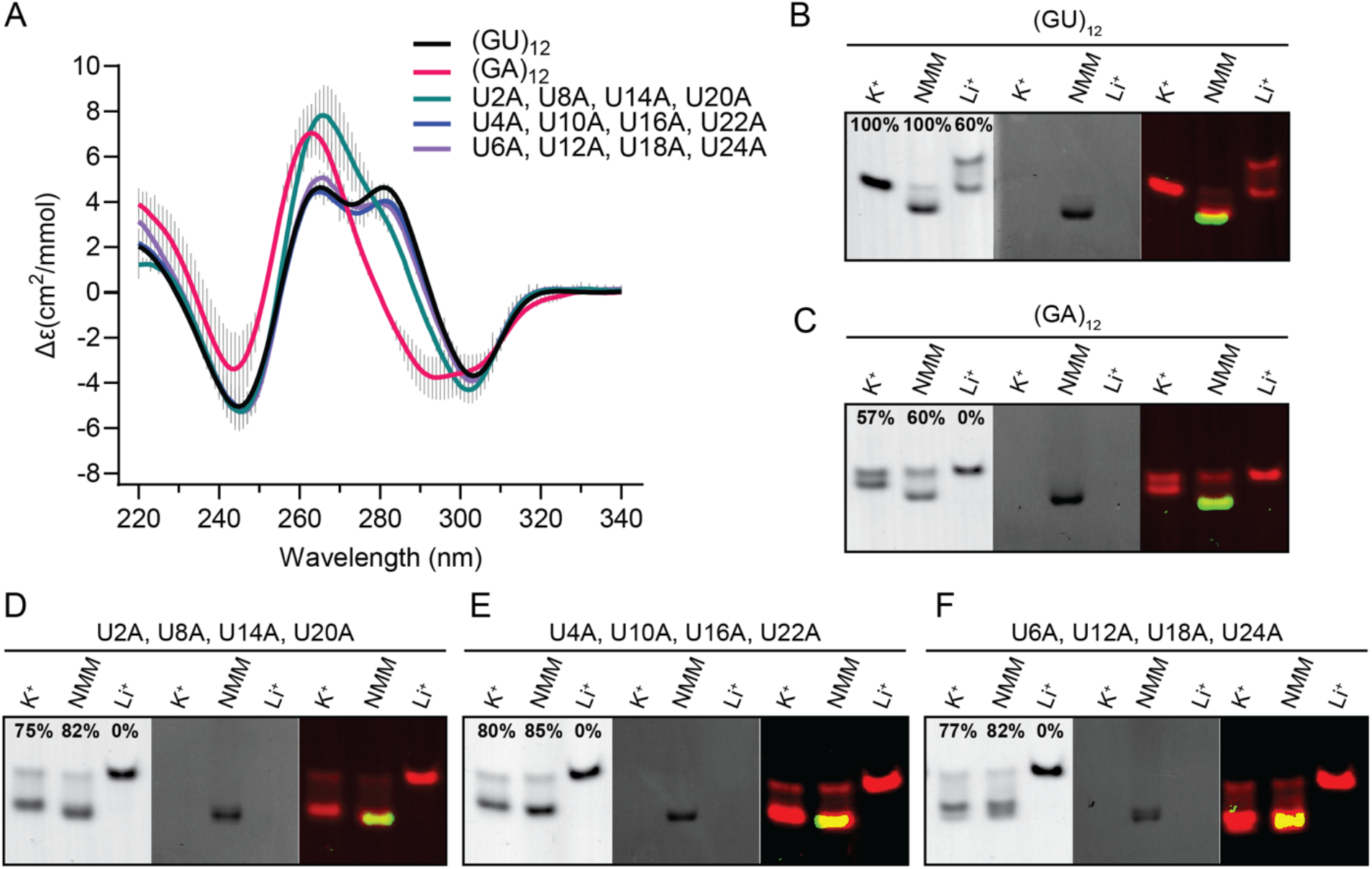
(A) CD spectra of pUG folds with multiple A substitutions including quadruple U to A variants and (GA)_12_. (B-F) Native gel electrophoresis of RNAs folded in K^+^ in the presence and absence of NMM, compared to unfolded RNA after Li^+^ treatment. RNAs were visualized using toluidine blue staining (left), NMM fluorescence (middle), and with both images overlayed in false colors (right). Data are shown for (B) (GU)_12_, (C) (GA)_12_, (D) U2A, U8A, U14A, U20A, (E) U4A, U10A, U16A, U22A and (F) U6A, U12A, U18A and U24A.

To further visualize the folding of these sequence variants, we analyzed them by EMSA and tested them for the ability to bind to *N*-methyl-mesoporphyrin (NMM), a G4 ligand that binds to the pUG fold with a *K*_D_ of 1 μM by stacking on the G1 quartet and forming hydrogen bonds with 2’ hydroxyl groups (1). When bound to NMM, the pUG fold migrates more quickly through the gel, likely due to the 2 negatively charged carboxylate groups on NMM (Figure 4B-F). All variants display potassium-dependent electrophoretic mobilities identical to the pUG fold and are unfolded when treated with lithium (Figure 4B-F). Additionally, all variants bind to NMM, consistent with G4 formation (Figure 4B-F). NMM fluorescence increases 60-fold when stacked upon G4s (1,24), and all variants showed this enhanced NMM fluorescence upon binding (Figure 4B-F). The (GA)_12_ RNA is folded as measured by CD (Figure 4A and Table 1) and is mostly folded in the EMSA assay (Figure 4C). The EMSA data show that the folded conformation of (GA)_12_ binds NMM, as expected for a G4 (Figure 4C). The quadruple U to A variants are well-folded in both the CD and EMSA assay and all bind NMM (Figure 4D-F and Table 1). We measured the thermodynamic stabilities of the quadruple U to A substitutions and the (GA)_12_ variant, which have low melting temperatures of 32-36 °C (Supplemental Figure 1 and Supplemental Table 1). Therefore, multiple U to A substitutions significantly destabilize the pUG fold. Nevertheless, all variants can adopt pUG-like folds which include the characteristic left-handed *syn-anti* stacking interactions within the center of the fold.

We next tested whether the pUG fold can tolerate multiple deoxyribose substitutions. Our previous NMR data showed that single deoxyribose substitutions are tolerated by the pUG fold at each of the 6 different positions in the first hexameric repeat (3). Four out of the six nucleotide conformations in the hexamer (positions 1, 2, 5 and 6) have 2’ endo sugar puckers, which is favored by deoxyribose (1,3). However, the 2’ hydroxyl groups of the second and third nucleotides form water-mediated hydrogen bonds to a partially hydrated K^+^ ion (1)(Figure 1A and B). We therefore hypothesized that the pUG fold may tolerate deoxyribose substitution at positions which are 2’ endo but do not form known ionic interactions, which corresponds to positions 1, 5 and 6 of each hexamer. Indeed, the d(G1, G5, T6)x4 variant is folded (Figure 5A), although not as well as (GU)_12_ (0.52 vs. 0.75, respectively) (Table 1). Deoxythymidine substitution is well tolerated at the second position, and the d(T2, T8, T14, T20) variant folds better than (GU)_12_ (Figure 5A and Table 1). The pUG fold cannot be entirely DNA, as neither d(GU)_12_ or d(GT)_12_ fold (Figure 5B). Some variants with 50-66% deoxyribose are partially folded (Figure 5A), while others are unfolded (Figure 5B). These data indicate that the pUG fold can tolerate partial, but not complete deoxyribose substitution.

**Figure 5.**
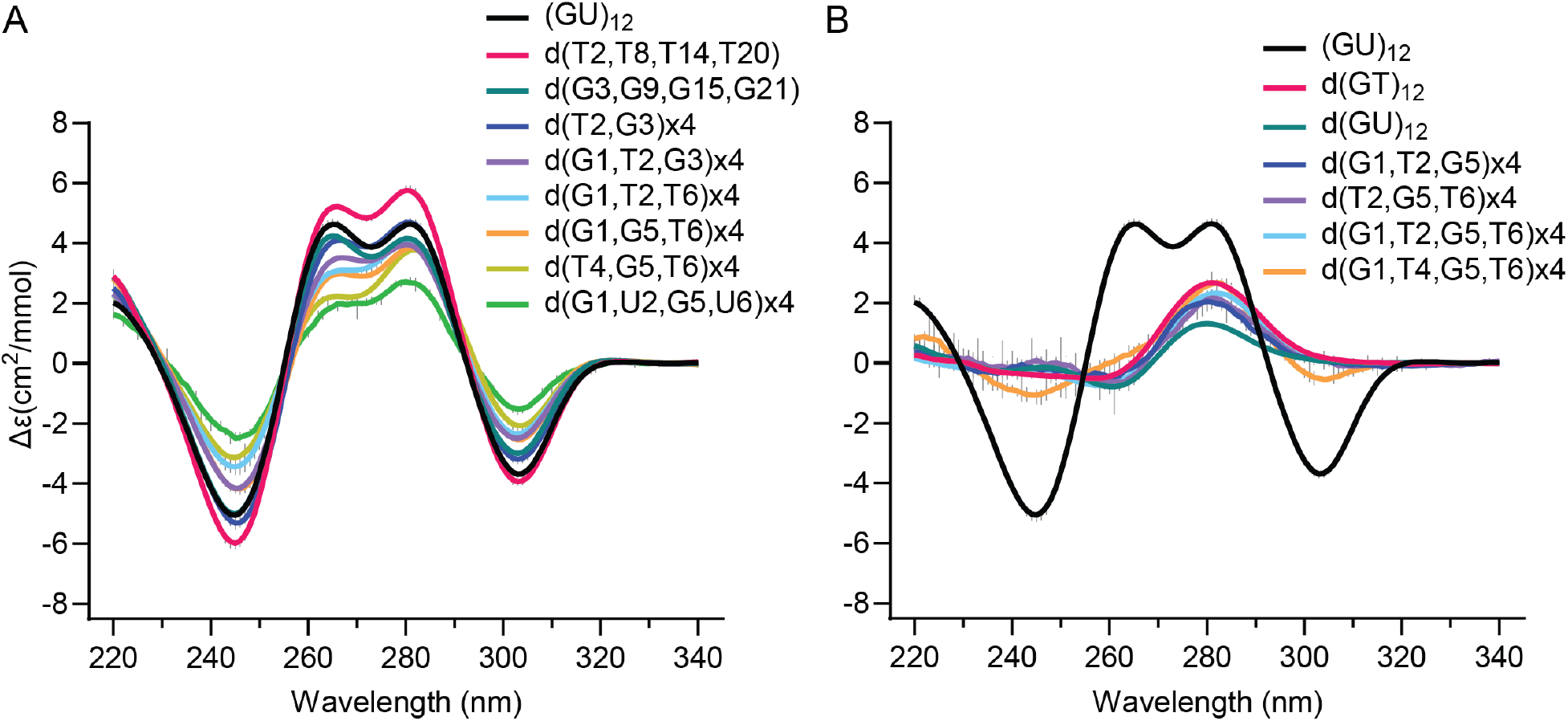
CD spectra of (A) deoxy substitutions that are folded and (B) deoxy substituted variants that are unfolded. For reference, the (GU)_12_ pUG fold is shown in black. Variants designated x4 include deoxy substitutions at all symmetry-related positions.

### Ionic requirements for pUG fold formation

We investigated the cation dependence of pUG folding in potassium, ammonium and sodium containing buffers. We find the pUG fold specifically requires potassium ions and cannot fold in sodium or ammonium (Figure 6A). We also investigated pUG folding in 150 mM KCl and 2 mM MgCl_2_, and in a pseudo-physiological buffer condition with mixed cations (140 mM KCl, 10 mM NaCl, 2 mM MgCl_2_, 0.3 mM spermine and 0.4 mM spermidine) (15) (Figure 6A). We titrated potassium and fit the resulting CD data to the Hill equation to determine the apparent equilibrium folding constant (K_0.5_ = 6.4 mM) and the Hill coefficient (n = 2.4) for potassium (Figure 6B and C). The Hill coefficient is consistent with cooperative uptake of multiple potassium ions, as observed in the structure. We performed CD temperature melting experiments to compare thermodynamic stabilities and observed little difference between the potassium only (T_m_= 52 °C) and potassium with magnesium (T_m_= 51 °C)(Supplemental Figure 2). These data indicate that magnesium does not stabilize the pUG fold. However, we observe a significant decrease in stability in the pseudo-physiological buffer (T_m_= 45 °C)(Supplemental Figure 2). This decrease in stability can be attributed to the presence of polyamines, which have been previously observed to destabilize quadruplexes with short loops (15).

**Figure 6.**
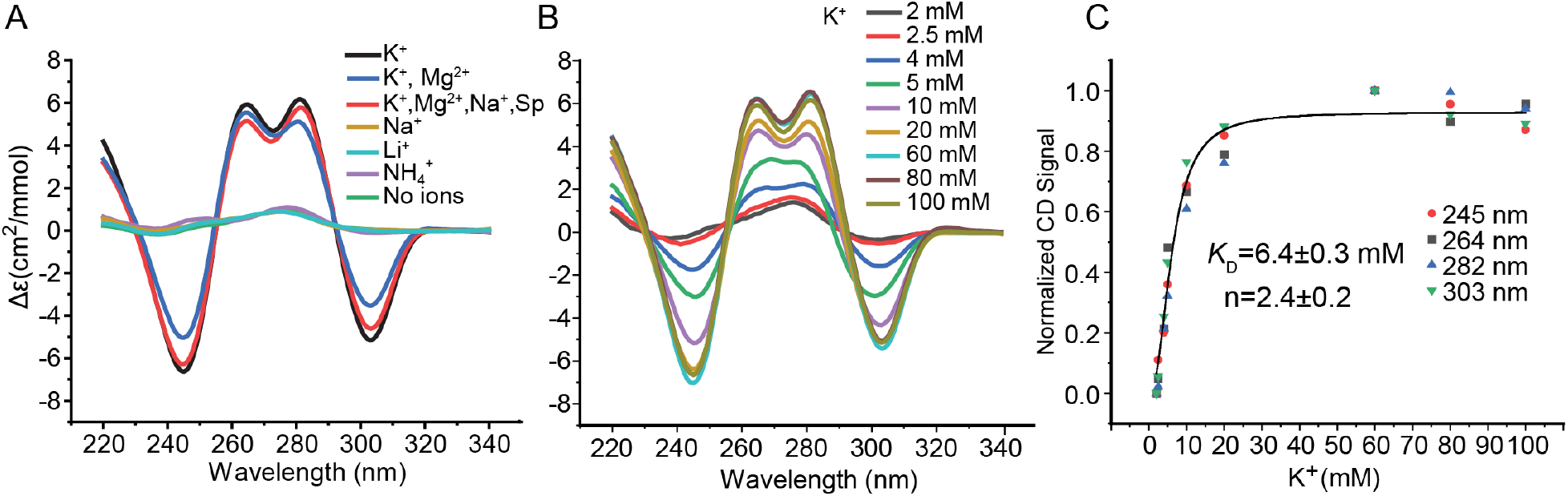
(A) (GU)_12_ RNA folding in buffers with different cations including: 150 mM K^+^, 150 mM K^+^ and 2 mM Mg^2+^, 140 mM K^+^, 10 mM Na^+^, 2 mM Mg^2+^, 0.3 mM spermine and 0.4 mM spermidine (K^+^+Mg^2+^+Na^+^+SP), 150 mM Na^+^, 150 mM Li^+^, 150 mM NH4^+^, and no ions (20 mM Tris buffer pH 7.0). (B) K^+^ concentration dependence of pUG folding monitored by CD at specified K^+^ concentrations. All samples were 20 μM RNA. (C) Normalized CD data from (B) fit to the Hill equation.

### Flanking sequences affect pUG folding

We previously showed that the pUG fold can form in the middle of an RNA when flanked by polyadenosines (1), and at RNA 3’ ends (3). To investigate the effect of structured flanking sequences on pUG folding, we created 4 different RNA constructs, each 47 nt in length. We reasoned that complementary flanking sequences might stabilize the pUG fold, similar to how G4-containing fluorogenic aptamers are stabilized by adjacent A-form helices (25-29). All sequences contained AA linkers between the pUG fold and the flanking sequences (Figure 7), similar to the fluorogenic aptamers (25-29). As a control, we created a construct with randomly generated 5’ and 3’ flanking sequences with little to no predicted base pairing between these flanking sequences (Figure 7A, random tails). Three additional constructs were generated with base pairing between the flanking sequences: one with AU base pairs (bps), one with mixed bps and one with bps based on the original mango aptamer sequence (25)(Figure 7A). All constructs folded except for the random tail sequence (Figure 7B). An increase in the intensity of the 260 nm peak is observed for the AU bps, mixed bps and mango bps sequences, consistent with the formation of pUG folds with additional A-form secondary structures. As the pUG fold is 24 out of 47 nucleotides, complete pUG fold formation as measured by the change in molar ellipticity at 304 nm should produce a fraction folded of ∼24/47, or 0.51. Analysis of the CD data for the AU bps, mixed bps and mango bps sequences show that indeed, these RNAs have folded fractions very close to 0.51 (Table 1). To further understand the folding of these RNAs, we performed RNA secondary structure predictions using energy minimization based on nearest-neighbor free energies (30). These predictions show the mixed bp and mango tail constructs have “unpaired” pUG regions that are available for pUG folding (Supplemental Figure 3). The secondary structure prediction for the AU bp construct pairs the pUG sequence in a way that precludes pUG folding, which is clearly incorrect and highlights inherent challenges with predicting quadruplexes (Supplemental Figure 3). However, the prediction for the random tail construct aligns with the CD data, as this RNA does not adopt a pUG fold. The random tail construct included the di- and trinucleotide sequences CA and CAC, which are perfectly complementary to pUGs, and allows formation of a stem-loop structure that likely prevents pUG folding (Supplemental Figure 3).

**Figure 7.**
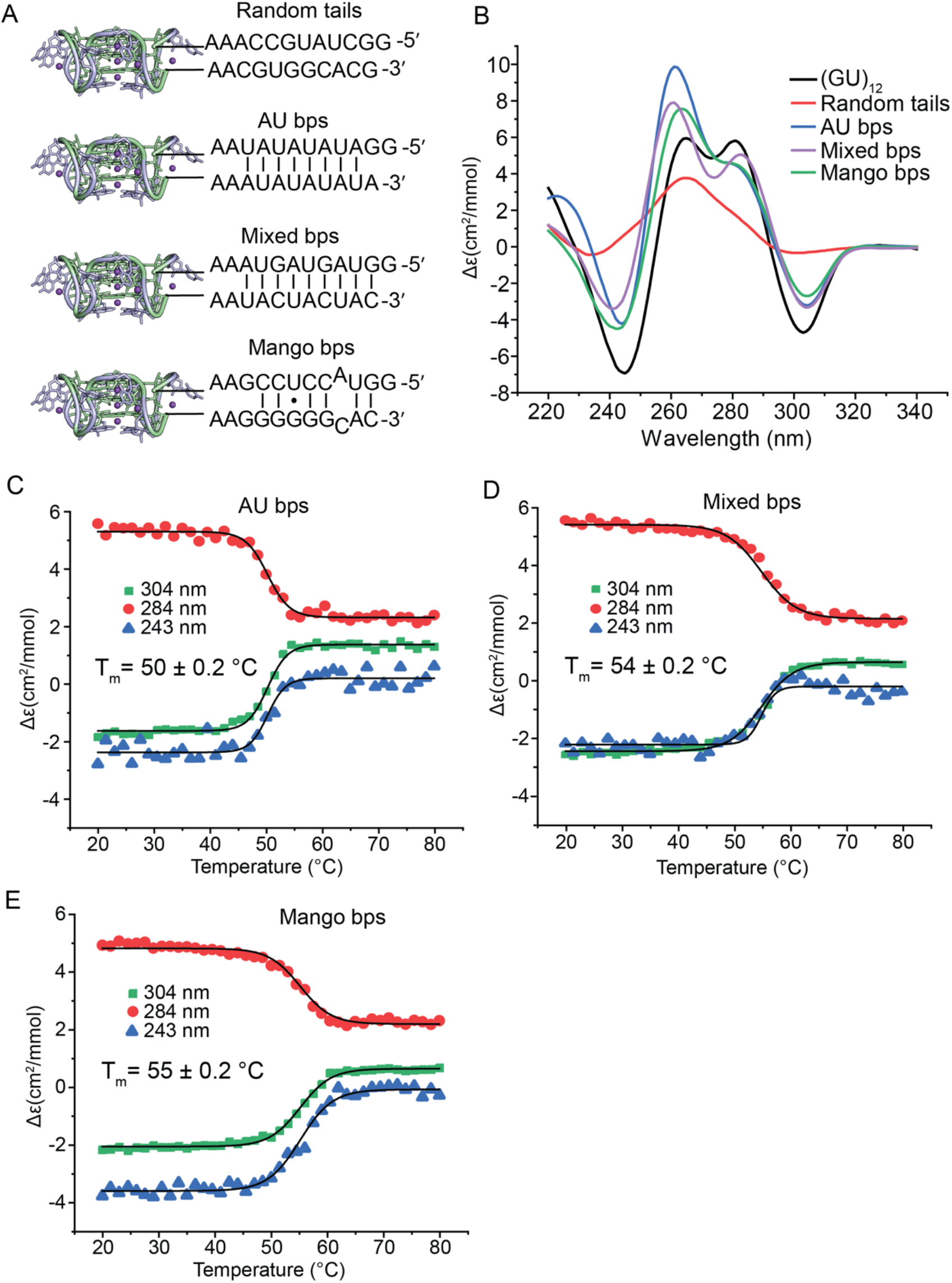
(A) 47 nucleotide RNAs with indicated 5’ and 3’ flanking sequences attached to (GU)_12_. (B) CD spectra of RNAs. (C-E) Thermal melts of (C) AU bps RNA, (D) mixed bps and (E) mango bps constructs.

We measured the thermodynamic stabilities of the pUG folds in the mango, AU bp and mixed bp tail constructs and compared them to the isolated (GU)_12_ pUG fold. The pUG folds in the mango tails and mixed bp tail RNAs are more stable than the isolated pUG fold (T_m_ = 54-55 vs 52 °C) while the AU-rich tail pUG fold was slightly destabilized (T_m_ = 50 °C)(Figure 7). The observation that the mango tails and mixed bp tails constructs are more thermodynamically stable than the isolated pUG fold is consistent with the flanking helices having a stabilizing effect on the pUG fold. In contrast, other flanking sequences can destabilize or interfere with folding, especially if they contain CA-rich regions that are complementary to pUGs. Thus, pUG fold formation and stability are highly dependent on sequence context.

## Discussion

Here we show that the pUG fold is not limited to poly(UG) RNA. We systematically tested a total of 40 sequence variants for pUG folding (Table 1). From these data, we ascertain that the pUG fold consensus sequence is flexible aside from a requirement for 12 guanosines. The spacing of the 12 guanosines is also flexible, as we have previously shown that the pUG fold can tolerate AA insertions (1). Our data indicate that any single uridine in the fold can be substituted by any other nucleotide. Single uridine substitutions and multiple U to A substitutions are well tolerated. U to G variants adopt pUG folds but show some degree of misfolding that can be attributed to the GGG stretches produced by these variants, which likely induce heterogenous G4 interactions. Additionally, multiple uridines cannot be substituted with cytidine, as this would create self-complementary GC-rich sequences that form stable hairpin or duplex structures. As few as 2 or 3 U to C substitutions, depending upon their position in the sequence, may result in alternative secondary structures with CG base pairs that prevent pUG folding. Indeed, formation of 3 C-G base pairs can effectively prevent pUG folding (Figure 7 and Supplemental Figure 2).

To our knowledge, this is the first report of GA repeat RNA forming left-handed G4 structures. GA repeat RNAs have been found to associate with replicated DNA (31). The idea that RNAs with GU and GA repeats may form similar G4 structures is consistent with previous work showing that both GU and GA repeats inhibit ribosome scanning and induce frameshifting, but only in the presence of G4-stabilizing ligands (32). While the (GA)_12_ structure forms a pUG-like fold at room temperature, its thermodynamic stability is significantly lower than the pUG fold (32 vs. 52 °C). We find that 4 U to A substitutions is enough to significantly decrease the stability of the pUG fold (T_m_ = 34-36 °C). Since the primary source of nucleic acid thermodynamic stability is base stacking, destabilization via adenosine substitution is likely due to adenosine stacking interactions that disrupt the fold. This is consistent with the observation that adenosine substitution at U2 and symmetry-related positions disrupts G1-G3 stacking, as evidenced by loss of the corresponding 280 nm CD peak (Figure 4A). In contrast, adenosine substitution at the U quartet, which is on a solvent exposed edge of the structure, preserves the complete CD spectrum but still destabilizes the fold, likely via disruption of K^+^ ion coordination that is maintained in part by the U quartet carbonyl oxygens (Figure 1A and B). Thus GA-rich sequences may require stabilization by ligands or additional folded elements to adopt pUG folds at physiological temperatures. Ligands that can stabilize pUG folds include small molecules such as porphyrins (1) and proteins. The human protein DNMT1 is one such pUG fold binding protein (11). While the pUG fold does not form in DNA, DNA GA repeats have been previously observed to form 4-stranded structures (33-35). DNA GT repeats have also been hypothesized to form G4 structures under conditions of extreme molecular crowding (36). However, we find no evidence for G4 formation by DNA GT repeats in the absence of molecular crowding.

The pUG fold has a high affinity and specificity for potassium ions, unlike other quadruplexes that have more relaxed ionic requirements and can form in potassium as well as sodium or ammonium (35,37-41). The pUG fold’s selectivity for potassium ions can be attributed to either the optimal fit of the ion, the greater energetic cost of sodium ion dehydration (42), or a combination of these effects. The K_0.5_ of 6 mM K^+^ and n = 2.4 values measured here are within the range of observed values for other G4 structures (43). Highlighting the pUG fold’s selectivity for K^+^ is the fact that its stability is unaffected by the addition of physiological concentrations of Mg^2+^, which is unusual for an RNA structure. This may be due to the fact that the pUG fold binds 6 potassium ions, an average of 1 cation for every 4 nucleotides (Figure 1) (1). Although Mg^2+^ does not affect the melting temperature of the pUG fold, it does accelerate its folding kinetics (20). The observation that the polyamines spermine and spermidine lower the T_m_ of the pUG fold is consistent with previous studies of other quadruplexes (15). Possibly, the linear array of cationic charges on spermine (+4) and spermidine (+3) may promote linear nucleic acid conformations (15) while destabilizing the pUG fold which has 8 backbone turns and inversions (Figure 1).

The data presented here expand our understanding of nucleic acid folding and demonstrate that diverse, commonly occurring RNA sequences can form pUG folds. Currently, it is not possible to predict pUG fold formation, as further improvements in RNA secondary and 3D structure prediction methods are needed and this remains a challenging area of computational biology. While the human transcriptome contains >20,000 perfect sequence matches to the (GU)_12_ pUG fold sequence (1), we find there are an additional ∼4100 pUG sequence variants with single U to N substitutions that can potentially adopt pUG folds (Supplemental Figure 4). The human transcriptome also contains ∼2400 GA repeats with at least 12 Gs (Supplemental Figure 4). Taken together as an upper limit, we estimate there are ∼27,000 sequences that could potentially adopt pUG folds within the human transcriptome, which is more than the number of human genes. However, we also show that pUG folds are highly sensitive to flanking sequences that can either stabilize or destabilize the fold. It therefore seems likely that a prerequisite for pUG fold formation is the absence of competing high probability secondary structures. On the other hand, pUG fold formation is favored when surrounding sequences are complementary. Base pairing of complementary sequences prevents competing interactions while simultaneously bringing the 5’ and 3’ ends into proximity, which is a requirement for the pUG fold (Figure 1A). In *C. elegans*, pUG tails are added to RNA 3’ ends and can be over 100 nt long (2). The formation of pUG folds in *C. elegans* is a requirement for the recruitment of RNA dependent RNA polymerase for the synthesis of secondary siRNAs (1). Long pUGs at 3’ ends likely provide an optimal context for pUG folding *in vivo*, as there are no neighboring sequences to interfere with folding. Given the large number of potential pUG fold sequences within genomes, it will be interesting in the future to determine when and where they form in other organisms.

## Methods

### RNA Production

RNAs were transcribed using T7 RNA polymerase. RNAs were purified using denaturing 15% polyacrylamide gel (7 M Urea, 100 mM Tris, 90 mM Boric acid, 10mM EDTA) electrophoresis. The RNA was identified using UV-shadowing, cut with a razor, and removed from the gel by diffusion at room temperature overnight in 300 mM sodium acetate, 50 mM HCl and 1 mM EDTA, pH 5.6, and then passed through a 0.2-μm filter. RNAs were further purified using a 1 ml Hi-trap Q column (GE Healthcare) that was equilibrated in 100 mM NaCl, 10 mM KH2PO4, 10 mM K2HPO4 and 1 mM EDTA. RNA was washed with 20 ml of buffer, then eluted with equilibration buffer with 2 M NaCl. RNAs were concentrated in an Amicon Ultra 3-kDa filter and buffer-exchanged into nuclease-free water (Invitrogen), to which the indicated buffer and salt concentrations were added. All RNA samples were pre-folded by addition of the 20 mM Bis-tris pH.7 and the appropriate salts, heating the samples in 1 l of 90 °C water and slowly cooling to room temperature for 5–6 h.

### RNA oligonucleotides used in this study

(GU)_12_: 5’-GUGUGUGUGUGUGUGUGUGUGUGU-3’

G7A: 5’-GUGUGUAUGUGUGUGUGUGUGUGU-3’

G7C: 5’-GUGUGUCUGUGUGUGUGUGUGUGU-3’

G7U: 5’-GUGUGUUUGUGUGUGUGUGUGUGU-3’

U8A: 5’-GUGUGUGAGUGUGUGUGUGUGUGU-3’

U8C: 5’-GUGUGUGCGUGUGUGUGUGUGUGU-3’

U8G: 5’-GUGUGUGGGUGUGUGUGUGUGUGU-3’

G9A: 5’-GUGUGUGUAUGUGUGUGUGUGUGU-3’

G9C: 5’-GUGUGUGUCUGUGUGUGUGUGUGU-3’

G9U: 5’-GUGUGUGUUUGUGUGUGUGUGUGU-3’

U10A: 5’-GUGUGUGUGAGUGUGUGUGUGUGU-3’

U10C: 5’-GUGUGUGUGCGUGUGUGUGUGUGU-3’

U10G: 5’-GUGUGUGUGGGUGUGUGUGUGUGU-3’

G11A: 5’-GUGUGUGUGUAUGUGUGUGUGUGU-3’

G11C: 5’-GUGUGUGUGUCUGUGUGUGUGUGU-3’

G11U: 5’-GUGUGUGUGUUUGUGUGUGUGUGU-3’

U12A: 5’-GUGUGUGUGUGAGUGUGUGUGUGU-3’

U12C: 5’-GUGUGUGUGUGCGUGUGUGUGUGU-3’

U12G: 5’-GUGUGUGUGUGGGUGUGUGUGUGU-3’

U2A: 5’-GAGUGUGUGUGUGUGUGUGUGUGU-3’

U4A: 5’-GUGAGUGUGUGUGUGUGUGUGUGU-3’

U6A: 5’-GUGUGAGUGUGUGUGUGUGUGUGU-3’

U2A, U8A, U14A, U20A: 5’-GUGAGUGAGUGUGAGUGUGAGUGU-3’

U4A, U10A, U16A, U22A: 5’-GUGAGUGUGAGUGUGAGUGUGAGU-3’

U6A, U12A, U18A, U24A: 5’-GUGUGAGUGUGAGUGUGAGUGUGA-3’

(GA)_12_: 5’-GAGAGAGAGAGAGAGAGAGAGAGA-3’

Random tails: 5’-GGCUAUGCCAAAGUGUGUGUGUGUGUGUGUGUGUGUAACGUGGCACG-3’

AU tails: 5’-GGAUAUAUAUAAGUGUGUGUGUGUGUGUGUGUGUGUAAAUAUAUAUA-3’

Mixed bp tails: 5’-GGUAGUAGUAAAGUGUGUGUGUGUGUGUGUGUGUGUAAUACUACUAC-3’

Mango tails: 5’-GGUACCUCCGAAGUGUGUGUGUGUGUGUGUGUGUGUAAGGGGGGCAC-3’

d(GT)_12_: 5’-dGdTdGdTdGdTdGdTdGdTdGdTdGdTdGdTdGdTdGdTdGdTdGdT-3’

d(GU)_12_: dGdUdGdUdGdUdGdUdGdUdGdUdGdUdGdUdGdUdGdUdGdUdGdU-3’

d(T2,T8,T14,T20): 5’-GdTGUGUdTGUGUGdTGUGUGdTGUGU-3’

d(G3,G9,G15,G21): 5’-GUdGUGUGUdGUGUGUdGUGUGUdGUGU-3’

d(T2,G3)x4: 5’-GdTdGUGUGdTdGUGUGdTdGUGUGdTdGUGU-3’

d(G1,T2,G3)x4: 5’-dGdTdGUGUdGdTdGUGUdGdTdGUGUdGdTdGUGU-3’

d(G1,T2,G5)x4: 5’-dGdTGUdGUdGdTGUdGUdGdTGUdGUdGdTGUdGU-3’

d(G1,T2,T6)x4: 5’-dGdTGUGdTdGdTGUGdTdGdTGUGdTdGdTGUGdT-3’

d(G1,G5,T6)x4: 5’-dGUGUdGdTdGUGUdGdTdGUGUdGdTdGUGUdGdT-3’

d(T2,G5,T6)x4: 5’-GdTGUdGdTGdTGUdGdTGdTGUdGdTGdTGUdGdT-3’

d(T4,G5,T6)x4: 5’-GUGdTdGdTGUGdTdGdTGUGdTdGdTGUGdTdGdT-3’

d(G1,T2,G5,T6)x4: 5’-dGdTGUdGdTdGdTGUdGdTdGdTGUdGdTdGdTGUdGdT-3’

d(G1,U2,G5,U6)x4: 5’-dGdUGUdGdUdGdUGUdGdUdGdUGUdGdUdGdUGUdGdU-3’

d(G1,T4,G5,T6)x4: 5’-dGUGdTdGdTdGUGdTdGdTdGUGdTdGdTdGUGdTdGdT-3’

### Native gel analyses of RNA folding

RNA samples were 10 μM. N-methyl mesoporphyrin IX (NMM) was added at 10 μM. RNA samples were mixed with an equal volume of 40% sucrose and loaded onto a 1 mm-thick native 10% polyacrylamide gel (29:1 acrylamide:bis acrylamide) containing KCl and Tris–borate–EDTA buffer (100 mM Tris, 90 mM Boric acid, 10mM EDTA, 5 mM KCl) and run at 5 W for 2 h at 4 °C. Gels were stained with 0.1% toluidine blue and imaged using a CanoScan LIDE300 image scanner (Canon). Fluorescent NMM samples were imaged with an Amersham Typhoon 5 gel scanner (Cytiva) with 488 and 670 nm wavelength filters for excitation and emission, respectively.

### Circular dichroism

CD samples contained 20 μM RNA in 20 mM Tris buffer pH 7.0 and either 150 mM KCl or 150 mM LiCl. For temperature melting experiments, the RNA samples contained 20 μM RNA in 20 mM potassium phosphate buffer pH 7.0 and 130 mM KCl. CD spectra were recorded in an AVIV model 420 CD spectrometer using a quartz cell with an optical path length of 1 mm. The scans were carried out with a step size of 1 nm and 5-s averaging times, and measurements were taken from 210 to 340 nm. Spectra were measured at 25 °C with buffer subtraction, and data were converted to molecular CD absorption, Δε:

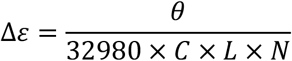

where θ is the raw CD signal (in millidegrees), C is the RNA concentration (in M), L is the cuvette path length (in cm), and N is the number of nucleotides. Thermal denaturation studies were also carried out by heating each sample from 20 to 85 °C in 1.5 °C intervals and with a 5-min equilibration time at each temperature. The ellipticity was measured at four different wavelengths (244, 264, 284 and 340 nm) with an averaging time of 10 s at each temperature. The thermal unfolding profile was characterized by the millidegree signal at 244, 264 and 284 nm to determine Tm and ΔG(37 °C) values by global fitting of the data to the Modified Gibbs-Helmholtz equations using Origin 2020 (OriginLab Corporation)(23). The potassium dependence of RNA folding was determined by collecting full wavelength scans as above. The 244, 264, 282, and 304 nm peaks were normalized with the maximum CD signal set to 1 to determine the equilibrium folding constant (K_0.5_) and hill coefficient (n) by globally fitting the data to the Hill equation using Origin (Origin 2020 OriginLab corporation).

## References

1. Roschdi, S., Yan, J., Nomura, Y., Escobar, C.A., Petersen, R.J., Bingman, C.A., Tonelli, M., Vivek, R., Montemayor, E.J., Wickens, M. et al. (2022) An atypical RNA quadruplex marks RNAs as vectors for gene silencing. Nature Structural & Molecular Biology, 29, 1113–1121.

2. Shukla, A., Yan, J., Pagano, D.J., Dodson, A.E., Fei, Y., Gorham, J., Seidman, J.G., Wickens, M. and Kennedy, S. (2020) poly(UG)-tailed RNAs in genome protection and epigenetic inheritance. Nature, 582, 283–288.

3. Escobar, C.A., Petersen, R.J., Tonelli, M., Fan, L., Henzler-Wildman, K.A. and Butcher, S.E. (2023) Solution Structure of Poly(UG) RNA. J Mol Biol, 435, 168340.

4. Butcher, S.E. (2024) A left-handed RNA quadruplex directs gene silencing. Trends Biochem Sci, 49, 387–390.

5. Kwon, D. (2025) RNA function follows form - why is it so hard to predict? Nature, 639, 1106–1108.

6. Abramson, J., Adler, J., Dunger, J., Evans, R., Green, T., Pritzel, A., Ronneberger, O., Willmore, L., Ballard, A.J., Bambrick, J. et al. (2024) Accurate structure prediction of biomolecular interactions with AlphaFold 3. Nature, 630, 493–500.

7. Degenhardt, M.F.S., Degenhardt, H.F., Bhandari, Y.R., Lee, Y.T., Ding, J., Yu, P., Heinz, W.F., Stagno, J.R., Schwieters, C.D., Watts, N.R. et al. (2025) Determining structures of RNA conformers using AFM and deep neural networks. Nature, 637, 1234–1243.

8. Geng, A., Roy, R. and Al-Hashimi, H.M. (2024) Conformational penalties: New insights into nucleic acid recognition. Curr Opin Struct Biol, 89, 102949.

9. Rodgers, M.L. and Woodson, S.A. (2019) Transcription Increases the Cooperativity of Ribonucleoprotein Assembly. Cell, 179, 1370–1381 e1312.

10. Lin, C.L., Taggart, A.J. and Fairbrother, W.G. (2016) RNA structure in splicing: An evolutionary perspective. RNA Biol, 13, 766–771.

11. Jansson-Fritzberg, L.I., Sousa, C.I., Smallegan, M.J., Song, J.J., Gooding, A.R., Kasinath, V., Rinn, J.L. and Cech, T.R. (2023) DNMT1 inhibition by pUG-fold quadruplex RNA. Rna, 29, 346–360.

12. Malgowska, M., Gudanis, D., Kierzek, R., Wyszko, E., Gabelica, V. and Gdaniec, Z. (2014) Distinctive structural motifs of RNA G-quadruplexes composed of AGG, CGG and UGG trinucleotide repeats. Nucleic Acids Research, 42, 10196–10207.

13. Guiset Miserachs, H., Donghi, D., Börner, R., Johannsen, S. and Sigel, R.K. (2016) Distinct differences in metal ion specificity of RNA and DNA G-quadruplexes. J Biol Inorg Chem, 21, 975–986.

14. Balaratnam, S. and Basu, S. (2015) Divalent cation-aided identification of physico-chemical properties of metal ions that stabilize RNA G-quadruplexes. Biopolymers, 103, 376–386.

15. Williams, A.M., Poudyal, R.R. and Bevilacqua, P.C. (2021) Long Tracts of Guanines Drive Aggregation of RNA G-Quadruplexes in the Presence of Spermine. Biochemistry, 60, 2715–2726.

16. Sun, H., Xiang, J., Liu, Y., Li, L., Li, Q., Xu, G. and Tang, Y. (2011) A stabilizing and denaturing dual-effect for natural polyamines interacting with G-quadruplexes depending on concentration. Biochimie, 93, 1351–1356.

17. Amarantos, I., Zarkadis, I.K. and Kalpaxis, D.L. (2002) The identification of spermine binding sites in 16S rRNA allows interpretation of the spermine effect on ribosomal 30S subunit functions. Nucleic Acids Res, 30, 2832–2843.

18. Ouameur, A.A., Bourassa, P. and Tajmir-Riahi, H.A. (2010) Probing tRNA interaction with biogenic polyamines. Rna, 16, 1968–1979.

19. Lightfoot, H.L., Hagen, T., Cléry, A., Allain, F.H. and Hall, J. (2018) Control of the polyamine biosynthesis pathway by G(2)-quadruplexes. Elife, 7.

20. Petersen, R.J., Vivek, R., Tonelli, M., Roschdi, S. and Butcher, S.E. (2025) The structure, folding kinetics, and dynamics of long poly(UG) RNA. Nucleic Acids Res, 53.

21. Banco, M.T. and Ferre-D’Amare, A.R. (2021) The emerging structural complexity of G-quadruplex RNAs. RNA, 27, 390–402.

22. Gajarsky, M., Stadlbauer, P., Sponer, J., Cucchiarini, A., Dobrovolna, M., Brazda, V., Mergny, J.L., Trantirek, L. and Lenarcic Zivkovic, M. (2024) DNA Quadruplex Structure with a Unique Cation Dependency. Angew Chem Int Ed Engl, 63, e202313226.

23. Greenfield, N.J. (2006) Using circular dichroism collected as a function of temperature to determine the thermodynamics of protein unfolding and binding interactions. Nat Protoc, 1, 2527–2535.

24. Sabharwal, N.C., Savikhin, V., Turek-Herman, J.R., Nicoludis, J.M., Szalai, V.A. and Yatsunyk, L.A. (2014) N-methylmesoporphyrin IX fluorescence as a reporter of strand orientation in guanine quadruplexes. FEBS J, 281, 1726–1737.

25. Dolgosheina, E.V., Jeng, S.C., Panchapakesan, S.S., Cojocaru, R., Chen, P.S., Wilson, P.D., Hawkins, N., Wiggins, P.A. and Unrau, P.J. (2014) RNA mango aptamer-fluorophore: a bright, high-affinity complex for RNA labeling and tracking. ACS Chem Biol, 9, 2412–2420.

26. Lu, X., Passalacqua, L.F.M., Nodwell, M., Kong, K.Y.S., Caballero-Garcia, G., Dolgosheina, E.V., Ferre-D’Amare, A.R., Britton, R. and Unrau, P.J. (2024) Symmetry breaking of fluorophore binding to a G-quadruplex generates an RNA aptamer with picomolar KD. Nucleic Acids Res, 52, 8039–8051.

27. Trachman, R.J., 3rd, Autour, A., Jeng, S.C.Y., Abdolahzadeh, A., Andreoni, A., Cojocaru, R., Garipov, R., Dolgosheina, E.V., Knutson, J.R., Ryckelynck, M. et al. (2019) Structure and functional reselection of the Mango-III fluorogenic RNA aptamer. Nat Chem Biol, 15, 472–479.

28. Trachman, R.J., 3rd, Demeshkina, N.A., Lau, M.W.L., Panchapakesan, S.S.S., Jeng, S.C.Y., Unrau, P.J. and Ferre-D’Amare, A.R. (2017) Structural basis for high-affinity fluorophore binding and activation by RNA Mango. Nat Chem Biol, 13, 807–813.

29. Yang, M., Prestwood, P.R., Passalacqua, L.F.M., Balaratnam, S., Fullenkamp, C.R., Arney, J.W., Weeks, K.M., Ferre-D’Amare, A. and Schneekloth, J.S., Jr. (2025) Structure-informed design of an ultrabright RNA-activated fluorophore. Nat Chem, 17, 1188–1195.

30. Mittal, A., Ali, S.E. and Mathews, D.H. (2024) Using the RNAstructure Software Package to Predict Conserved RNA Structures. Curr Protoc, 4, e70054.

31. Gylling, H.M., Gonzalez-Aguilera, C., Smith, M.A., Kaczorowski, D.C., Groth, A. and Lund, A.H. (2020) Repeat RNAs associate with replication forks and post-replicative DNA. RNA, 26, 1104–1117.

32. Morren, B.M., Marcelis, J., Muradin, I. and Olsthoorn, R.C.L. (2025) Investigation of possible G-quadruplex formation by GU- and GA-rich repeats and their role in translation. RNA Biol, 22, 1–12.

33. Lee, J.S., Evans, D.H. and Morgan, A.R. (1980) Polypurine DNAs and RNAs form secondary structures which may be tetra-stranded. Nucleic Acids Res, 8, 4305–4320.

34. Lee, J.S. (1990) The stability of polypurine tetraplexes in the presence of mono- and divalent cations. Nucleic Acids Res, 18, 6057–6060.

35. Novotny, A., Novotny, J., Kejnovska, I., Vorlickova, M., Fiala, R. and Marek, R. (2021) Revealing structural peculiarities of homopurine GA repetition stuck by i-motif clip. Nucleic Acids Res, 49, 11425–11437.

36. Trizna, L., Osif, B. and Viglasky, V. (2023) G-QINDER Tool: Bioinformatically Predicted Formation of Different Four-Stranded DNA Motifs from (GT)(n) and (GA)(n) Repeats. Int J Mol Sci, 24.

37. Dingley, A.J., Peterson, R.D., Grzesiek, S. and Feigon, J. (2005) Characterization of the cation and temperature dependence of DNA quadruplex hydrogen bond properties using high-resolution NMR. J Am Chem Soc, 127, 14466–14472.

38. Lim, K.W., Ng, V.C., Martin-Pintado, N., Heddi, B. and Phan, A.T. (2013) Structure of the human telomere in Na+ solution: an antiparallel (2+2) G-quadruplex scaffold reveals additional diversity. Nucleic Acids Res, 41, 10556–10562.

39. Scaria, P.V., Shire, S.J. and Shafer, R.H. (1992) Quadruplex structure of d(G3T4G3) stabilized by K+ or Na+ is an asymmetric hairpin dimer. Proc Natl Acad Sci U S A, 89, 10336–10340.

40. Schultze, P., Hud, N.V., Smith, F.W. and Feigon, J. (1999) The effect of sodium, potassium and ammonium ions on the conformation of the dimeric quadruplex formed by the Oxytricha nova telomere repeat oligonucleotide d(G(4)T(4)G(4)). Nucleic Acids Res, 27, 3018–3028.

41. Hud, N.V., Schultze, P., Sklenar, V. and Feigon, J. (1999) Binding sites and dynamics of ammonium ions in a telomere repeat DNA quadruplex. J Mol Biol, 285, 233–243.

42. Hud, N.V., Smith, F.W., Anet, F.A. and Feigon, J. (1996) The selectivity for K+ versus Na+ in DNA quadruplexes is dominated by relative free energies of hydration: a thermodynamic analysis by 1H NMR. Biochemistry, 35, 15383–15390.

43. Zhang, A.Y.Q. and Balasubramanian, S. (2012) The Kinetics and Folding Pathways of Intramolecular G-Quadruplex Nucleic Acids. Journal of the American Chemical Society, 134, 19297–19308.

