## Supplemental Data for "Sequence and ionic requirements of pUG fold quadruplexes"

Supplemental Table 1

| Sequence | $T_m$ °C |
|---|---|
| $(GU)_{12}$ (150 mM $K^+$) | 52 |
| $(GU)_{12}$ (150 mM $K^+$, 2 mM $Mg^{2+}$) | 51 |
| $(GU)_{12}$ ($K^+$, $Na^+$, $Mg^{2+}$, Sp) | 45 |
| U2A | 49 |
| U4A | 44 |
| U6A | 49 |
| U2A, U8A, U14A, U20A | 36 |
| U4A, U10A, U16A, U22A | 35 |
| U6A, U12A, U18A, U24A | 34 |
| $(GA)_{12}$ | 32 |
| AU bps | 50 |
| Mixed bps | 54 |
| Mango bps | 55 |

Supplemental Figure 1

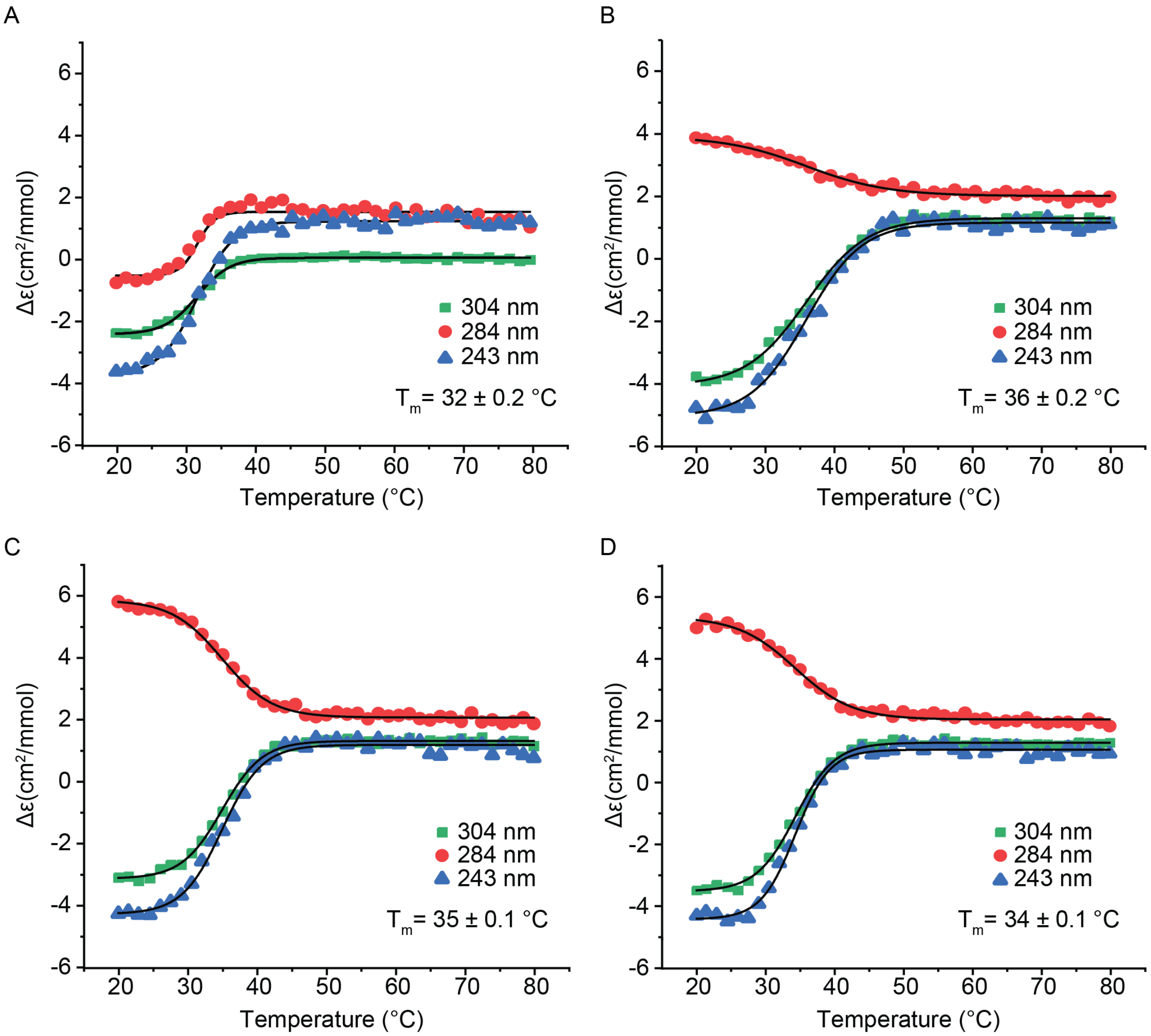


Supplemental Figure 1. CD monitored thermal denaturation of (A) (GA)12, (B) U2A, U8A, U14A, U20A, (C) U4A, U10A, U16A, U22A and (D) U6A, U12A, U18A and U24A. All data were fit to the Boltzmann equation to determine the $T_m$.

Supplemental Figure 2

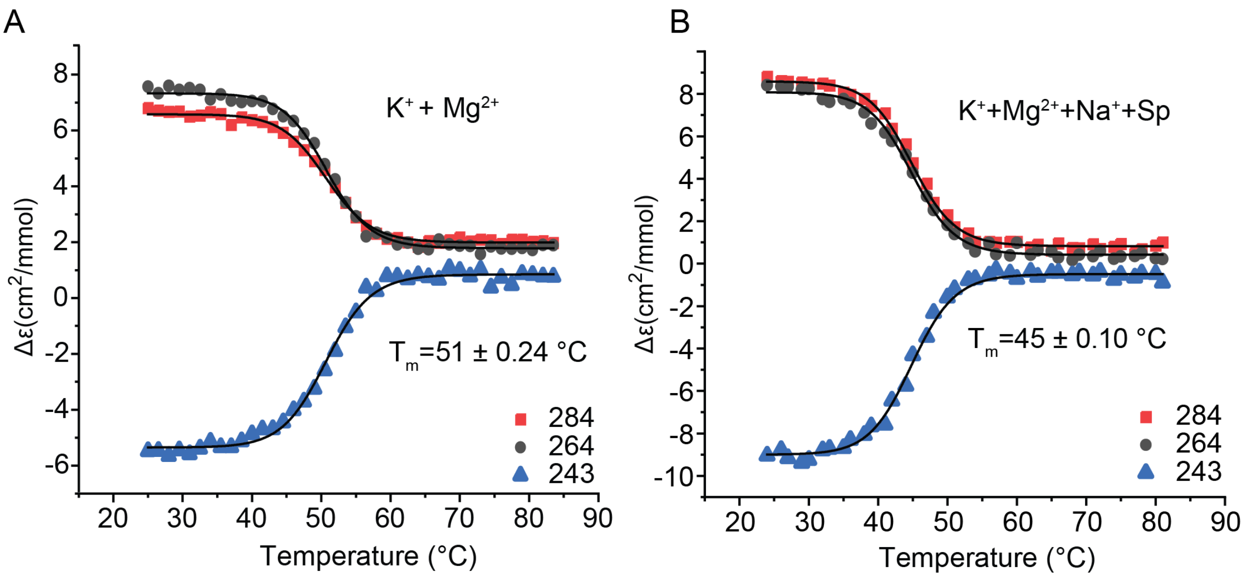


Supplemental Figure 2. (B) Thermal melt of $(GU)_{12}$ in 150 mM $K^+$ and 2 mM $Mg^{2+}$. (C) Thermal melt of $(GU)_{12}$ in $K^+$, $Na^+$, $Mg^{2+}$, Sp buffer (140 mM KCl, 10 mM NaCl, 2 mM $MgCl_2$, 0.3 mM spermine and 0.4 mM spermidine).

Supplemental Figure 3

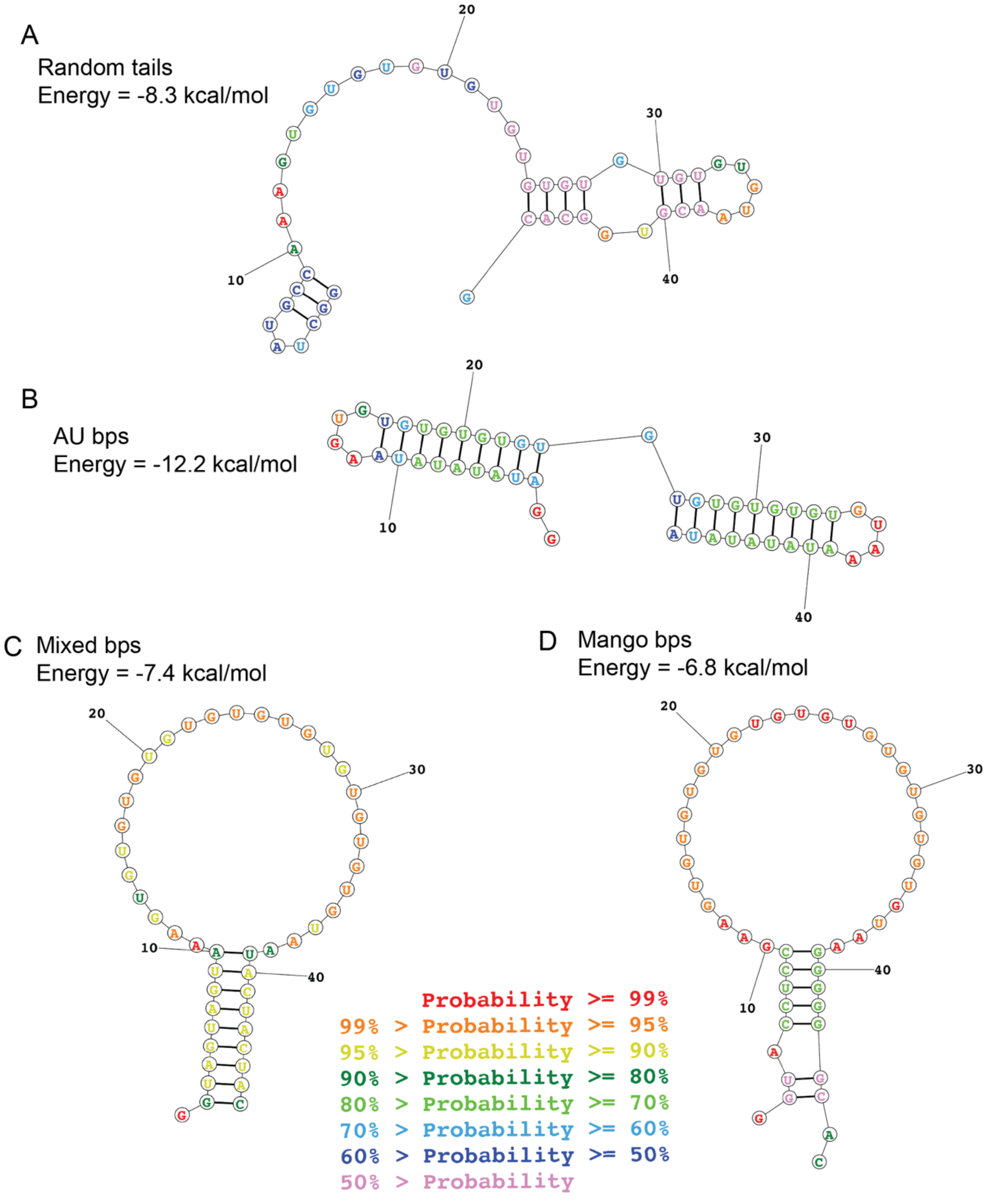


Supplemental Figure 3. Predicted secondary structure free energies and base pairing probabilities for 47 nucleotide RNAs with the indicated 5′ and 3′sequences flanking $(GU)_{12}$.

Supplemental Figure 4

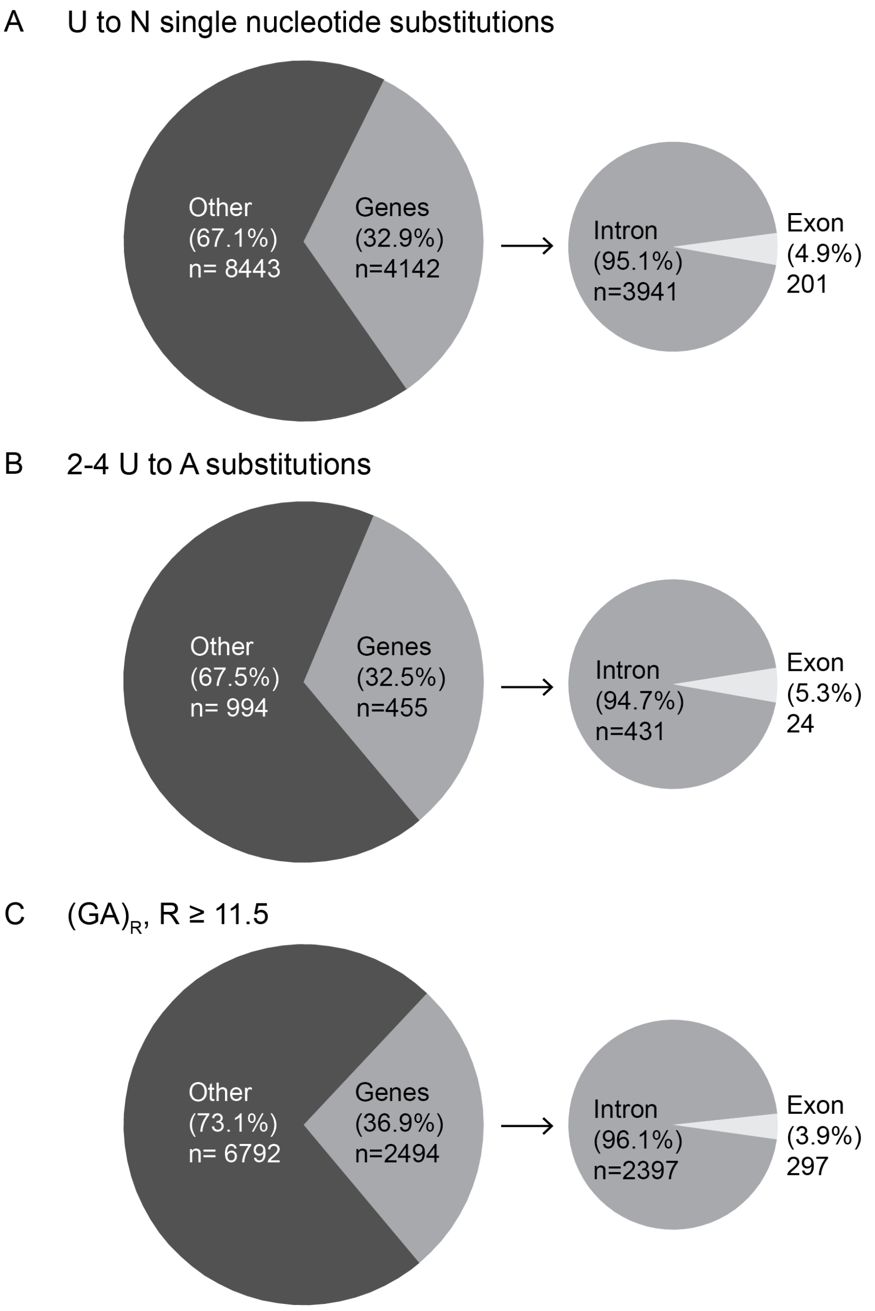


Supplemental Figure 4. Number and distribution of potential pUG fold sequences containing (A) single nucleotide U to N (where N is any nucleotide) substitutions, (B) 2-4 U to N substitutions, and (C) 11.5 or more GA repeats in the human genome.
